# Identification of further variation at the lipooligosaccharide outer core locus in *Acinetobacter baumannii* genomes and extension of the OCL reference sequence database for *Kaptive*

**DOI:** 10.1101/2023.02.23.529771

**Authors:** Bianca Sorbello, Sarah M. Cahill, Johanna J. Kenyon

## Abstract

The outer core locus (OCL) that includes genes for the synthesis of the variable outer core region of the lipooligosaccharide (LOS) is one of the key epidemiological markers used for tracing the spread of *Acinetobacter baumannii*, a bacterial pathogen of global concern. In this study, we screened 12476 publicly available *A. baumannii* genome assemblies for novel OCL sequences, detecting six new OCL types that were designated OCL17-OCL22. These were compiled with previously characterised OCL to create an updated version of the *A. baumannii* OCL reference database, providing a total of 22 OCL reference sequences for use with the bioinformatics tool, *Kaptive*. Use of this database against the 12476 downloaded assemblies found OCL1 to be the most common locus, present in 73.6% of sequenced genomes assigned by *Kaptive* with a match confidence score of ‘Good’ or above. OCL1 was most common amongst isolates belonging to sequence types, ST1, ST2, ST3 and ST78, all of which are over-represented clonal lineages associated with extensive antibiotic resistance. The highest level of diversity in OCL types was found in ST2, with eight different OCL identified. The updated OCL reference database is available for download from GitHub (https://github.com/katholt/Kaptive; under version *v 2.0.5*), and has been integrated for use on *Kaptive*-Web (https://kaptive-web.erc.monash.edu/) and PathogenWatch (https://pathogen.watch/), enhancing current methods for *A. baumannii* strain identification, classification and surveillance.

**IMPACT STATEMENT:** In the absence of effective treatment options for multi-drug resistant *Acinetobacter baumannii*, the highest-ranking critical priority bacterial pathogen of global concern, national and global surveillance is necessary to detect, track and subsequently curb the spread of isolates that resist current therapies. Several epidemiological markers are used to characterise *A. baumannii* strains by detecting genetic differences in specific regions of the genome. One of these is the chromosomal OC locus (OCL) responsible for the synthesis of the outer core (OC) component of the lipooligosaccharide (LOS). Here, we provide an update to the international *A. baumannii* OCL reference sequence database, extending the number of known OCL types to assist with clinical surveillance of important strains or clonal lineages.

**Data summary:**

1. The updated *A. baumannii* OCL reference sequence database including 22 annotated OCL sequences is available for download under *Kaptive v. 2.0.5* at https://github.com/katholt/Kaptive.
2. Genome assemblies or GenBank records used as representative reference sequences are listed in Table 1 and acknowledged in each record in the database.

## INTRODUCTION

*Acinetobacter baumannii* is a Gram-negative coccobacillus that is recognised as one of six leading pathogens responsible for nearly three quarters of deaths associated with antibiotic resistance worldwide [1]. The species has been detected in most geographical regions around the world and has been found to account for more than one fifth of all hospital-acquired infections in Europe, the Eastern Mediterranean and Africa [2]. The World Health Organisation has ranked carbapenem-resistant *A. baumannii* as a ‘Priority 1: CRITICAL’ bacterial pathogen [3], and with an estimated 80% of circulating isolates now resistant to last line carbapenems [1], new therapeutic approaches are urgently needed. However, in the current absence of widely accessible, approved and effective treatments, local and international surveillance initiatives are required to control the continued spread of pan-resistant *A. baumannii* isolates.

Whole genome sequencing (WGS) represents a readily accessible standard for tracking the spread and evolution of resistance in *A. baumannii.* As the majority of carbapenem resistant isolates belong to two globally disseminated clonal complexes, known as Global Clone 1 (GC1) and Global Clone 2 (GC2), multi-locus sequence typing (MLST) is commonly used in the primary stages of strain characterisation to identify the clonal lineage. Two MLST schemes, Institut Pasteur (IP) and Oxford (OX), are available for *A. baumannii*, and GC1 and GC2 include predominately sequence type (ST) 1 and ST2 in the IP scheme [4]. However, as clones continue to evolve and separate into distinct sublineages with different resistance profiles [5–9], two additional epidemiological markers can be used in combination with MLST to further discriminate isolates. These are the K locus (KL) for synthesis of the capsular polysaccharide (CPS), and the outer core locus (OCL) for synthesis of the outer-core (OC) of the lipooligosaccharide (LOS) [10–13]. Both CPS and LOS are important cell-surface structures and virulence determinants for *A. baumannii*, and differences in the genetic content at the chromosomal K and OC loci can lead to structural changes in these complex surface molecules [10, 14, 15].

To simplify the ability to detect genetic differences at these loci, a user-friendly nomenclature system that assigns a KL or OCL number to any new combination of genes found at these loci was developed [10]. In 2020, the many different clusters of genes identified at these locations were compiled into KL and OCL reference databases compatible with the bioinformatics search tool, *Kaptive* [11]. This tool assigns a best match locus type to queried genome assemblies by screening individual genomes against these databases [16]. The first release of these databases included 92 KL and 12 OCL types, which are available in *Kaptive* versions *0.0.7-2.0.0* [11]. We recently performed a major update to the *A. baumannii* KL reference sequence database releasing a further 149 KL in version *2.0.1* [12]. However, while a further four OCL types (OCL13-OCL16) have been characterised and reported since the release of the original databases [17, 18], the OCL reference sequence database has not been updated.

Unlike many Gram-negative bacterial pathogens, *A. baumannii* does not produce a structurally variable O-antigen polysaccharide attached to the OC of the LOS by a WaaL ligase [10, 13, 14] to form an extended structure known as lipopolysaccharide (LPS). Structural variation is therefore predominately observed in the OC component, which is directed by differences in the genetic content at the OC locus [13,14]. The OC locus is located between conserved *ilvE* and *aspS* genes in the *A. baumannii* chromosome [10], and interruption of genes at this location has been shown to result in truncations to the core component of the LOS structure [14]. All OCL characterised to date include predominately genes that encode glycosyltransferase (*gtr*) enzymes for linking sugars together to grow the oligosaccharide OC structure, though some OCL include additional genes for sugar biosynthesis and/or modification [13]. Previously, it was found that the first 12 OCL types identified could be separated into two major genetic arrangements, referred to as Group A and Group B, respectively based on the presence of either a *pda1* or *pda2* gene near the start of the locus (Figure 1). These genes are in ‘Region 1’, which also includes a few further genes that are shared by all OCL sequences that fall into each group. Diversity in gene content that distinguishes OCL types has so far always been observed ‘Region 2’ (Figure 1).

**Figure 1.**
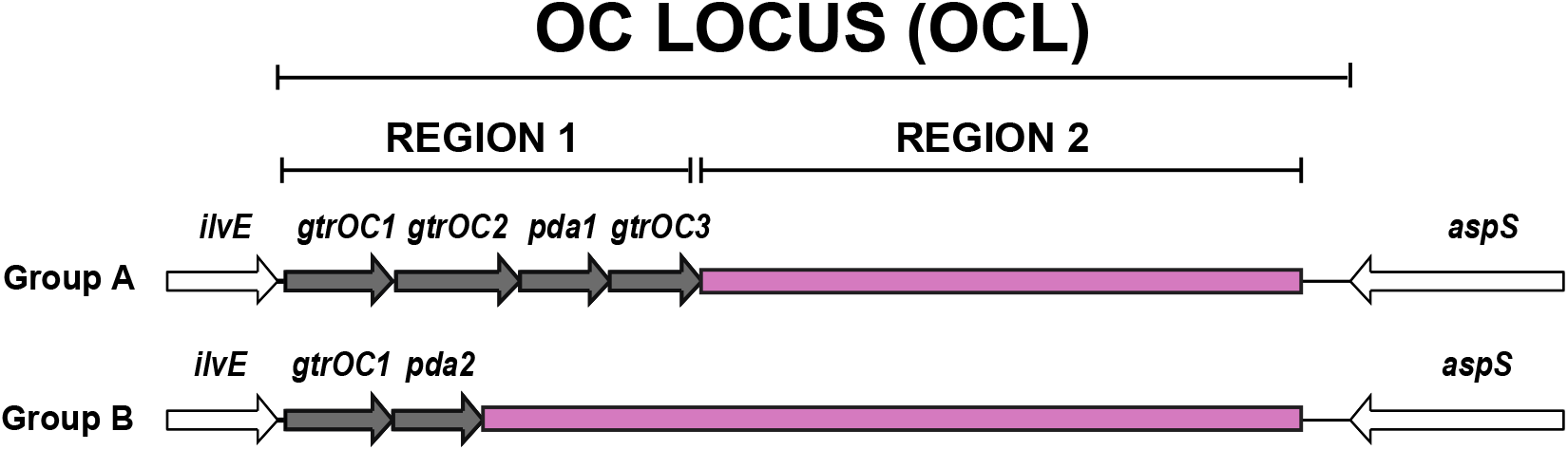
Typical arrangement of gene clusters belonging to Group A and Group B located at the *A. baumannii* chromosomal OC locus for lipooligosaccharide outer core biosynthesis. Conserved *ilvE* and *aspS* genes that flank the OC locus are shown in white. Grey genes are those common to OCL belonging to either Group A or Group B, whereas the pink segments are regions that commonly vary in gene content between different OCL types.

Here, we provide an update to the *A. baumannii* OCL reference sequence database to include the four additional OCL not included in the original version, and new OCL types identified among more than twelve thousand publicly available *A. baumannii* genome sequences. Additionally, we define the general characteristics of OCL sequences and assess the distribution of OCL types amongst sequenced isolates, revealing the extent of OCL diversity in clinically important clonal lineages.

## METHODS

### *A. baumannii* genome assemblies

Publicly available genome assemblies (n=12553) listed under the *Acinetobacter baumannii* taxonomical classification (ID: 470) in the NCBI non-redundant and WGS databases were downloaded for local analysis (21^st^ of April 2022). To confirm the taxonomical assignment, BLASTn was used to screen all downloaded genome assemblies for the presence of the *A. baumannii-specific oxaAb* gene (also known as *bla*_OXA-51-like_) as described previously [11, 12]. Only genomes confirmed to carry this gene, as defined by >90% combined coverage and >95% nucleotide sequence identity to the query *oxaAb* gene from *A. baumannii* isolate A1 (GenBank accession number CP010781.1, base positions 1753305 to 1754129) were used for further analyses.

### Detection and annotation of novel OC locus sequences in *A. baumannii* genomes

Using the command-line version of *Kaptive v. 2.0.0* [19], confirmed *A. baumannii* genomes (n=12476) were screened against an in-house version of the OCL database, which included OCL1-OCL12 reference sequences available in the published database (released under *Kaptive* versions *0.0.7-2.0.4;* https://github.com/katholt/Kaptive) plus the additional four OCL sequences (OCL13-OCL16) identified since its release. A “minimum gene identity” cut-off of 85% amino acid (aa) sequence identity was used as a parameter for *Kaptive* searches, which is the accepted cut-off for identification and annotation of OCL genes in *A. baumannii* [10, 11]. Matches with <95% total coverage and/or <95% total nucleotide sequence identity to the best matched sequence, additional or missing genes, and/or length discrepancies were selected for detailed manual inspection as described previously [11, 12].

OCL sequences were visually compared to their best match locus using the pairwise sequence comparison tool, EasyFig *v. 2.2.5* [20], using a tBLASTx identity cut-off of 85%. If differences were detected in gene presence/absence to the best match locus, the OCL was considered novel and assigned a new number according to the Kenyon and Hall nomenclature system for OC locus typing [10, 11]. In cases where OCL were found to include novel genes that share <85% aa sequence identity with known *A. baumannii* OCL gene products, BLASTp was used to search for gene homologues of known or predicted function and a new name was assigned to indicate enzyme function (e.g., GtrOC# for predicted glycosyltransferases) based on detection of protein motifs/domains using HMMER with the Pfam database [21].

### Creation and validation of an updated *A. baumannii* OCL reference sequence database

A reference GenBank format (.gbk) file was generated for each novel OCL sequence (details for each record are shown in Table 1), which included the complete nucleotide sequence of the locus and annotations of all coding sequences within that were assigned using the nomenclature scheme [10]. As only one OC structure, OC1, is known for *A. baumannii* [22], an additional note field was added to the OCL1 record to define the OC type as is required for the newest iteration of the *Kaptive* code [19]. For all other OCL where no OC structural data is available, the note field specifies the structural type as unknown.

**Table 1.**
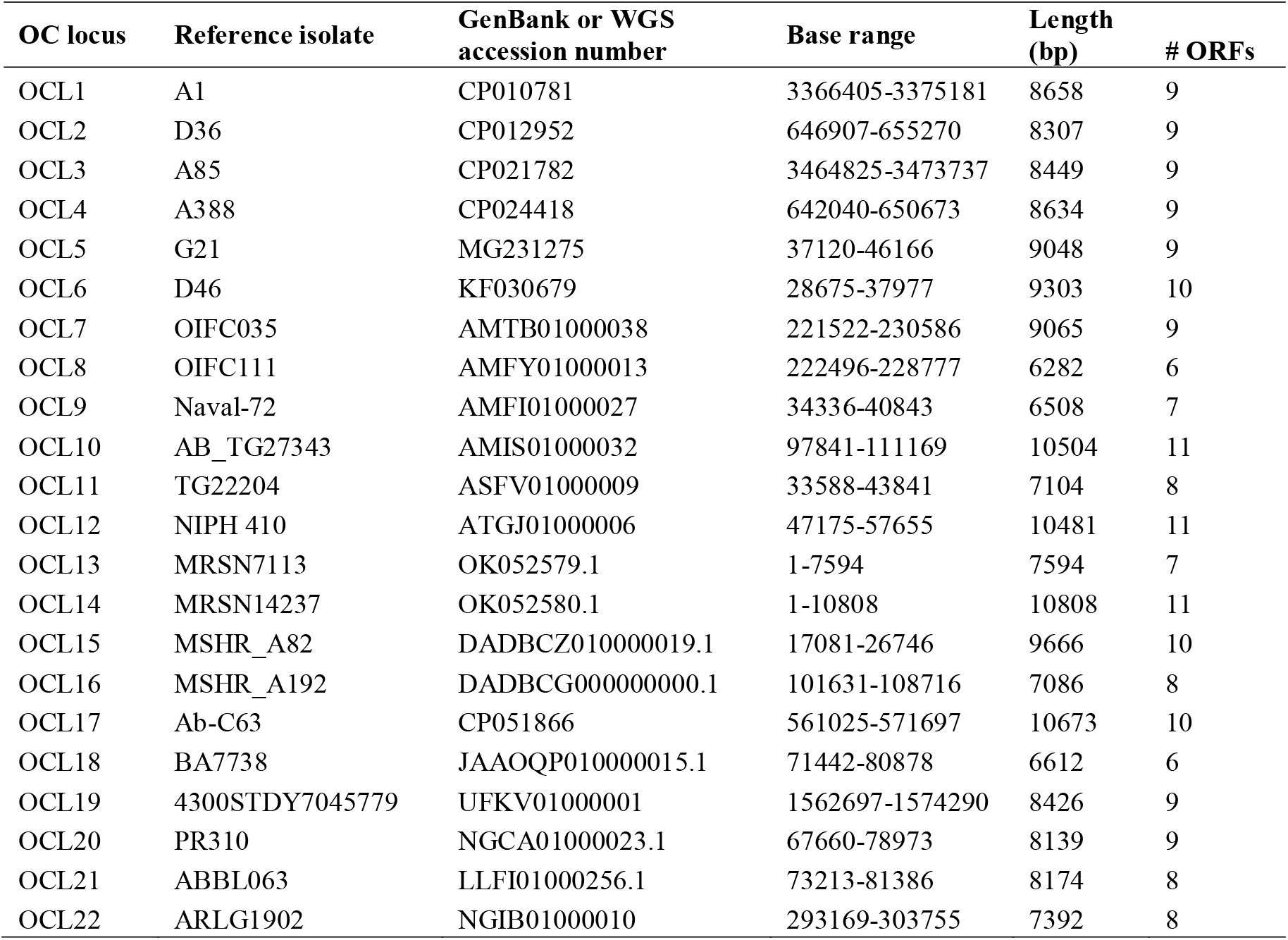
Information on *A. baumannii* OCL reference sequences used to populate the updated OCL database for *Kaptive v 2.0.5*

Reference .gbk files for all OCL were then concatenated to create an updated OCL database compatible with command-line *Kaptive*, which was released at https://github.com/katholt/Kaptive under version *v 2.0.5.* The updated database was also integrated with the *Kaptive*-Web (https://kaptive-web.erc.monash.edu/) and PathogenWatch (https://pathogen.watch/) platforms. Over the course of this analysis, a problem was identified that lead to the confusion of assignments for the closely related OCL5 and OCL13, and OCL1 and OCL18 loci. The problem was rectified through an update to the *Kaptive* code, which was released with the updated OCL database in *version 2.0.5.* To validate database functionality, all genome assemblies were searched again using *Kaptive v. 2.0.5* with the same parameters as described above.

### Characterisation of the genetic repertoire of *A. baumannii* OCL sequences

Prokka *v. 1.13* [23] was used to generate .gff3 files for each OCL reference sequence by directing the designation of gene names to annotations available in each .gbk record in the database. The .gff3 files were used to manually tabulate the sequence lengths and number of open reading frames (ORFs) for each OCL type. A gene presence/absence matrix based on a minimum gene identity cut-off of 85% aa sequence identity was also created using the pan genome analysis software, Roary *v. 3.13.0* [24]. Tabulated data was then visualised using the ggplot2 package in RStudio *v. 3.3.0+* [25]. A representative sequence from each gene homology group was submitted to HMMER [21] to re-assess protein family designations for the reported annotations. To further confirm that a gene for a potential WaaL ligase was not located between *aspS* and *tonB* in *A. baumannii*, all genomes were also screened for an insertion of additional sequence present at this location.

### Clonal analysis

Multi-locus sequence typing (MLST) was conducted on genome assemblies to assign sequence type (ST) using the MLST tool available at https://github.com/tseemann/mlst with the *A. baumannii* IP scheme. ST groups that included ≥100 genome representatives were identified as over-represented clonal groups, and the percentage of OCL types with a match confidence of ‘Good’ or above within each group was calculated and visualised as a heatmap using Rstudio *v. 3.3.0+* with the ggplot2 *v. 3.3.6* package [25].

## RESULTS

### Screening for *A. baumannii* genome assemblies that carry novel OCL

Confirmed *A. baumannii g*enome assemblies (n=12476) were screened against an in-house version of the OCL reference database that includes the 12 OCL available in *Kaptive v 0.7.0-2.0.0* plus the additional OCL13-OCL16 sequences identified since its release. To identify genomes that may carry novel OCL types, the 308 genome assemblies that were assigned a match confidence level of ‘Perfect’, indicating that the detected locus is identical to a known OCL reference sequence [26], were disregarded. An additional 2834 genomes (22.7% of the total genome pool) were also not further examined as the detected OCL was identified across two or more contigs, suggesting either low sequence quality or insertion sequence (IS) interruptions of OCL types as previously described [11, 12]. Manual inspection of all remaining genome assemblies (n=9334) revealed that 24 genomes could further be considered poor quality as the OCL included multiple gaps (>60 bp), insertions/deletions, or strings of ‘N’ bases. In addition, two other genomes (NCBI assembly accession numbers GCA_001862305.1 and GCA_001863295.1) had been assigned a best match locus with <19% coverage and 0% nucleotide sequence identity, and close inspection of these sequences revealed that an OC locus was not present.

A total of 9166 genomes (73.5% of the total genome pool) carried OC loci that were very close relatives of their best match locus, differing only in single nucleotide polymorphisms (SNPs) or small insertions/deletions. A further 104 genomes were considered variants of known loci as these were found to carry one or more IS interrupting the locus. For nine of these, the IS were found as part of one of two different novel transposons inserted at different locations in the OCL1 locus (shown in Supplementary Figure S1). One of these transposons was identified in two different locations, interrupting either *orf1(ghy)* in seven genomes or *gtr6* in a single genome. The other transposon was found in just one genome (NCBI assembly number GCA_021569095.1), interrupting the *gtrOC7* gene. For both transposons, the predicted gene products appear unlikely to influence OC biosynthesis. However, while additional sequence was detected between *ilvE* and *aspS*, these variants were not assigned new OCL names in accordance with the updated guidelines for KL/OCL nomenclature [12].

Amongst the remaining 38 genomes, six novel OCL, designated OCL17-OCL22, were identified. Of the novel types, OCL19 (one genome), OCL20 (one genome) and OCL22 (three genomes), included *pda1* and were classified as Group A, and OCL17 (26 genomes) and OCL21 (five genomes) included *pda2* and hence were classified as Group B (Figure 2). The OCL18 locus (present in two genomes) did not include either *pda1* or *pda2*, though it was found to be a deletion variant of OCL1 missing the *pda1, gtrOC3* and *gtrOC4* genes, hence it was therefore designated as Group A. Only four new genes were found amongst OCL20, OCL21 or OCL22, with three predicted to encode glycosyltransferases (named *gtrOC34-36)* and one (named *ORF4)* predicting a protein of unknown function (see below).

**Figure 2.**
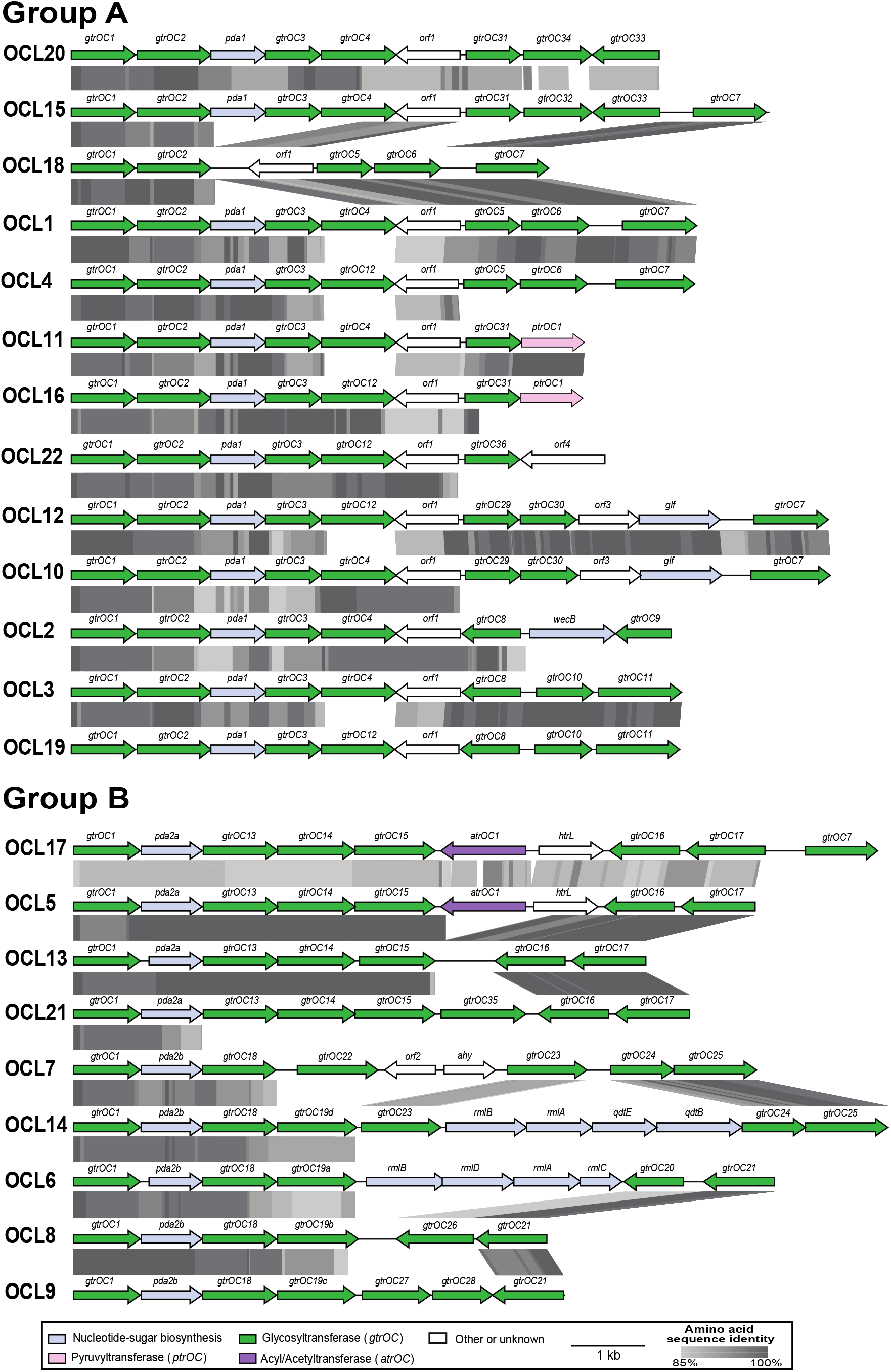
Alignment of all identified OCL reference sequences (OCL1–OCL22) separated into Group A and Group B. Horizontal arrows represent genes indicating their direction of transcription that are coloured by the predicted function of their gene products (legend shown below). Grey shading between gene clusters shows amino acid sequence identities determined by tblastx with grey scale shown in the key below. Figures drawn to scale using Easyfig [20] and annotated/coloured in Adobe Illustrator.

### Validation of the updated OCL reference sequence database

The six newly identified OCL types were added into an updated reference database, which included concatenated .gbk reference files for all known OCL types (OCL1-OCL22). To validate its use, the updated database was applied to the complete pool of 12476 genomes. Using the output of this search (Supplementary Table S1), the frequency of each OCL was calculated amongst the genomes with an assigned match confidence score of ‘Good’ or above (n=11940). OCL1 was found to be the most common type (Figure 3A), accounting for 73.6% (n=8782) of the genome pool. A further 23.4% of genomes (n=2797) included 1 of 6 other OCL types (OCL3 (8.05%), OCL6 (4.78%), OCL2 (4.28%), OCL5 (2.4%), OCL10 (2.09%), and OCL7 (1.83%)), and together with OCL1, represent 97.15% of the genomes analysed. The remaining 3% of genomes (n =361) contained one of the other 15 OCL types, each detected at a frequency <1%.

**Figure 3.**
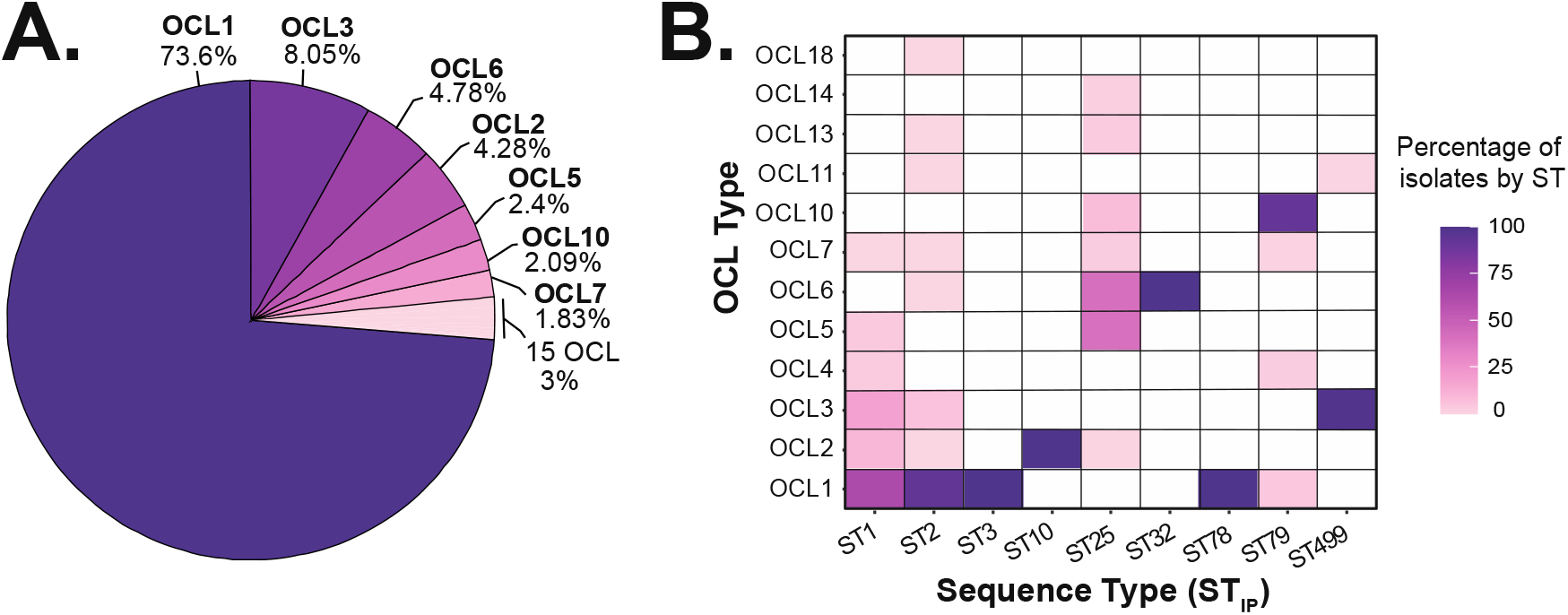
Distribution of OCL types amongst *A. baumannii* genome assemblies. **(A)** Percentage of OCL best match locus types assigned a confidence score of ‘Good’ or above by *Kaptive* (n = 11940). **(B)** Heat map showing percentage of genomes belonging to nine over-represented sequence types (STs) that carry OCL best match locus types assigned a confidence score of ‘good’ or above by *Kaptive.* Only STs with ≥100 genomes are shown (total number of genomes = 9331: 489 ST1, 7540 ST2, 254 ST3, 169 ST10, 221 ST25, 125 ST32, 109 ST78, 137 ST79, 287 ST499). Figures were created using ggplot2 package in RStudio [25].

### OCL in clonal genomes

The distribution of OCL amongst genomes from clonal isolates was also investigated. STs represented by more than 100 genome assemblies were identified, revealing nine clonal lineages (ST1, ST2, ST3, ST10, ST25, ST32, ST78, ST79 and ST499) that were over-represented in this genome pool. ST2 represented 63.15% (n=7540) of genomes, followed by ST1 representing 4.1% (n=489) of genomes. A total of 1302 genomes fell into the seven other ST groups, all of which include isolates from various geographical regions and associated with extensive antibiotic resistance. The calculated percentage of clonal isolates that carry each of the OCL types (Figure 3B), revealed that the predominant ST1 and ST2 lineages include 6 and 8 different OCL types, repectively, with OCL1 being the most common in both clones. The emerging, globally distributed clonal lineage, ST25, demonstrated a similar level of diversity at the OC locus, with 7 OCL types detected, for which OCL5 and OCL6 were the most common types. Only one OCL was detected in ST10, ST3, ST32 and ST78 lineages. Of the novel OCL sequences, OCL18 was identified at a low frequency in ST2, while OCL13 and OCL14 were detected in ST25 only.

### General characteristics of sequences at the OC locus

To better understand the diversity at the OC locus, the general features of all OCL types including the average length, number of ORFs, and frequency of OCL genes present, were investigated. OCL sequences were found to range between 6 kb and 10 kb (average 8 kb) in total length (Figure 4A; Table 1) and include between 6 and 11 (average 9) ORFs per locus (Figure 4B; Table 1). A gene presence/absence matrix categorising all ORFs across 22 OCL into ‘homology groups’ based on an 85% aa identity cut-off was generated using Roary to calculate how many ORFs were shared amongst OC locus types. A total of 57 genes (homology groups) were detected across the 22 OCL, and of these, 40.3% were found to be unique to a single locus type (Figure 4C). Only 1.8% of genes were found to be present in more than 75% of all OCL types, with *gtrOC1* being the only gene present in all loci.

**Figure 4.**
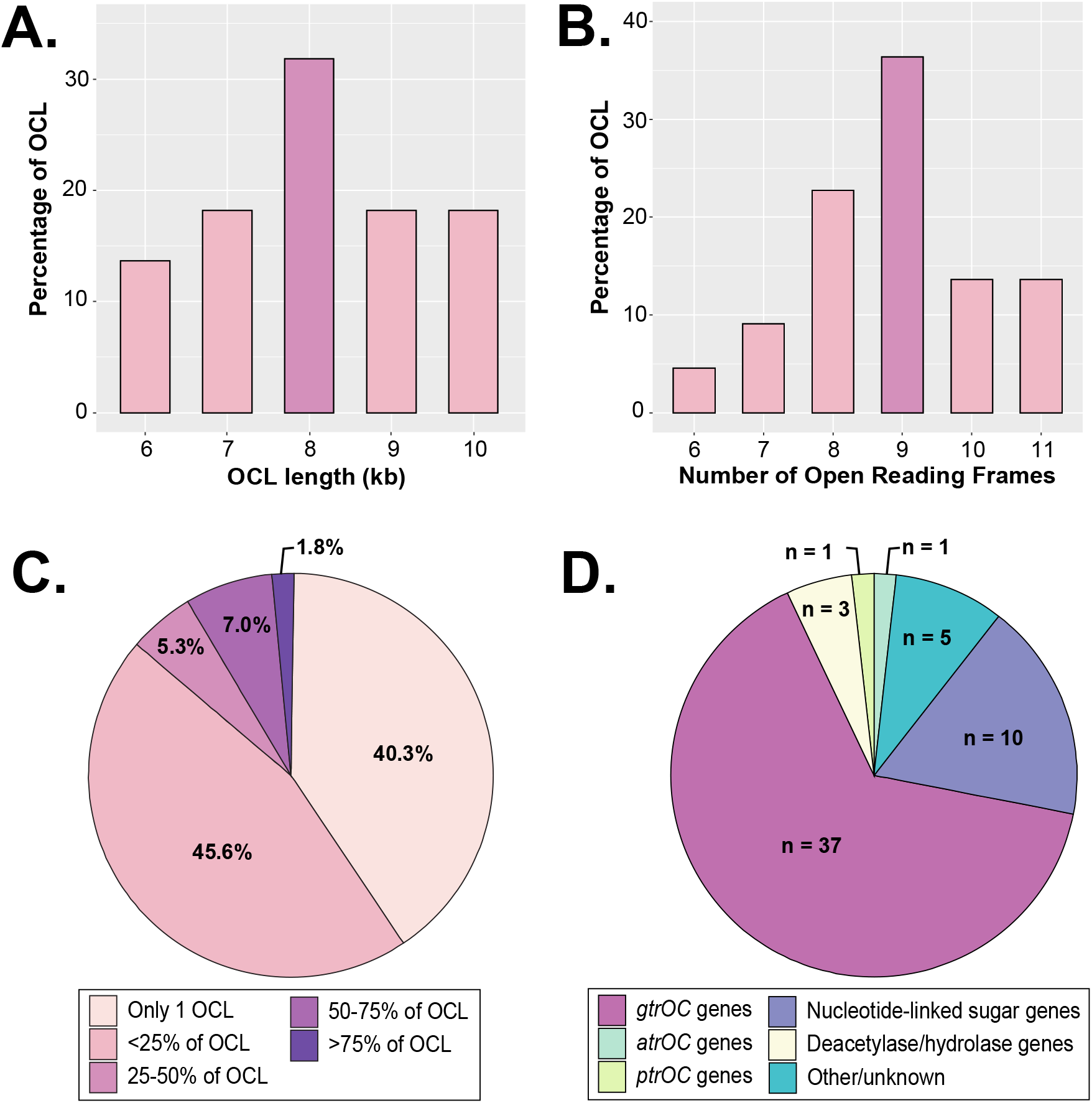
General features of the 22 OC locus sequences included in the database. **(A)** Percentage of OCL per total sequence length (kb). **(B)** Percentage of OCL that include specified number of ORFs. **(C)** Frequency of 57 gene ‘homology groups’ found across 22 OCL as determined by Roary gene presence/absence analysis. Colour scheme is shown below. **(D)** Breakdown of gene types found across 22 OCL at the OC locus. Figures were created using ggplot2 package in RStudio [25].

As the annotations of most genes included in the OCL reference database were originally assigned based on protein family (Pfam) and clan assignments from searches conducted in 2014 [13], the translated products of each homology group were subjected to HMMER to re-assess these assignments (Table 2). Of the 57 OCL genes, 37 were found to predict glycosyltransferases (GtrOC). The majority of these belong to glycosyltransferase-associated Pfams, confirming their designation as GtrOC proteins. However, consistent with previous searches, five proteins (GtrOC1, GtrOC14, GtrOC19a, GtrOC19b, and GtrOC22) were found to belong to the Mito_fiss_Elm1 (PF06258.14) mitochondrial fission protein family that has no known function in bacteria. One GtrOC protein (GtrOC7), not previously assigned to any family, was found to include domains for Stealth_CR1-CR4 families associated with capsular polysaccharide phosphotransferases. Two further proteins (GtrOC6 and GtrOC28) could not be assigned to a defined protein family, though GtrOC28 was previously found to belong to the Glyco_tranf_2_5 glycosyltransferase family in the previous study[13].

**Table 2.**
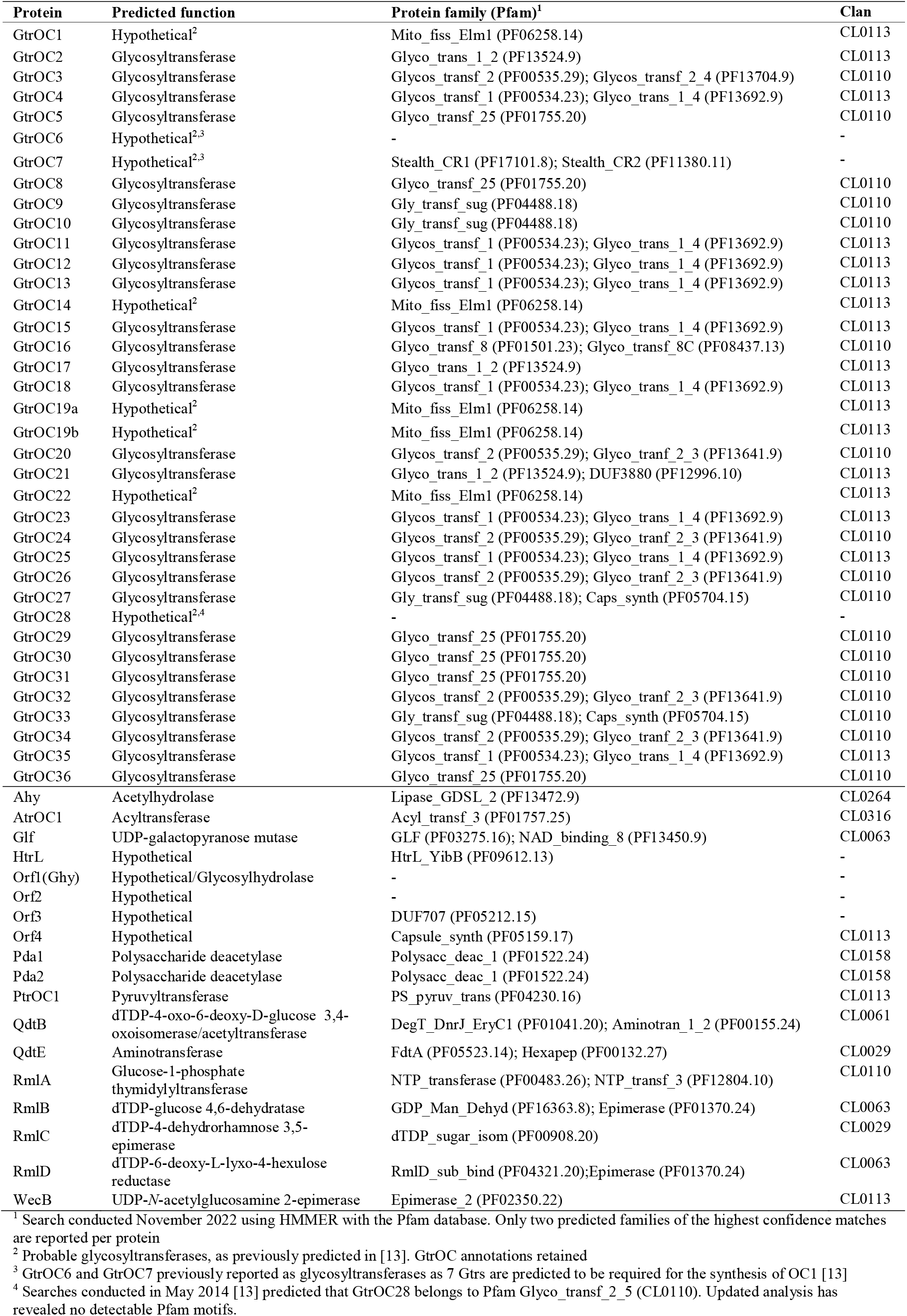
Predicted function of proteins encoded by OC locus genes

In addition to the 37 genes predicting GtrOC proteins, 10 genes were found to encode enzymes predicted to be responsible for nucleotide-linked sugar biosynthesis (either *rmlBDAC* for dTDP-L-rhamose, *rmlBA/qdtBE* for 3-acetamido-3,6-dideoxy-D-galactose, *wecB* for *N*-acetyl-D-mannosamine, and *glf* for D-galactofuranose). Two further genes are for predicted polysaccharide deacetylases (Pda1 and Pda2), one for a predicted acylhydrolase (Ahy), one for a putative acetyltransferase (AtrOC1), one for a putative pyruvyltransferase (PtrOC1), and five predicting proteins of unknown function (ORF1-4 and HtrL) (Table 2, Figure 4D). The *orf1* gene, found in all OCL in Group A, was originally predicted to encode a glycosylhydrolase and annotated as *ghy* in the earliest study on the OC locus [10]. However, this was since reassigned as *orf1* based on updated Pfam searches [13].

## DISCUSSION

In this study, we report an update to the *A. baumannii* OCL reference sequence database to include four OCL types (OCL13-OCL16) characterised since the release of the original database [17, 18] and six new OCL (OCL17-OCL22) identified amongst 9334 genomes, providing a new total of 22 distinct OCL types. While distribution of OCL types amongst sequenced genomes demonstrated that the most common OCL types had been accounted for in the original version of the database [11], the updated version encompasses a further ten OCL, increasing its utility for detecting a broader variety of OCL types in *A. baumannii.*

While new OCL were uncovered in this study, we cannot exclude the possibility that further novel types are present in the 2834 genomes that were not investigated due to the locus being detected across >1 contig. The finding of the OC locus in two or more contigs is often due to the occurrence of insertion sequences interrupting the locus [11, 12], and several previous studies have demonstrated that IS interruption of OCL is common, particularly in the over-represented OCL1 locus [8, 10, 13, 14]. The presence of a composite transposon interrupting OCL1 has also been described previously [8], and here we report further interruptions of OCL1 by two novel transposon sequences. It is not clear what properties are conferred by these transposons or the effect that these might have on the LOS. However, as the interruption of specific genes in OCL1 has been shown to lead to significant truncation of the LOS structure [14], the characterisation of the specific site of insertion(s) of either IS or transposons at the OC locus is recommended when assessing LOS structure or virulence properties associated with LOS in *A. baumannii.*

All novel types identified in this study resembled known OCL and could be classified into Group A or Group B, the two major OC locus configurations previously defined by the presence of common genes adjacent to *ilvE* [13] (Figure 1). Though no new variations to the general arrangement of the OC locus were uncovered, two subgroups within Group B could be observed, distinguished by the presence of either *gtrOC13* or *gtrOC18* immediately adjacent to *pda2.* Within Group A, the newly identified OCL18 locus was the only type found to vary in the genetic content of ‘Region 1’. OCL18 does not include *pda1* or *gtrOC3* but was assigned to Group A as it carries *gtrOC1, gtrOC2*, and *orf1* like other locus types in this group. Apart from OCL18 that may represent a subgroup within Group A, all other variation between OCL types was observed in ‘Region 2’ (Figure 1).

While most OCL are close relatives of each other, differing by single gene replacements or by the presence of a few additional genes (Figure 2), OCL18 and OCL13 are deletion variants of OCL1 and OCL5, respectively. The close relationship between OCL5 and OCL13 had been observed in a previous study [27], where OCL13 was found to lack the *atrOC1* gene and a complete copy of the adjacent *htrL* gene that are present in OCL5. Upon alignment of OCL5 and OCL13, we identified a 4 bp repeat (GTAA) in OCL5 on either side of the deleted region in OCL13, and a single copy of this 4 bp string in OCL13 at the precise location of the deletion (see Supplementary Figure S2A). Similarly, we identified a 4 bp repeat (CTAG) in OCL1 on either side of the deleted region, with a single copy found in OCL18 (Supplementary Figure S2B). This finding may suggest how one type arose from another, though further work is needed to confirm the effect on the LOS structure.

The genetic repertoire of the OC locus was found to be significantly less diverse than what has been observed at the *A. baumannii* K locus, which includes a repertoire of 681 genes across 237 KL types, for which 272 genes are predicted to encode glycosyltransferases [12]. However, like the K locus, the OC locus is considered a recombination ‘hotspot’ [7, 28], and most OCL genes are either unique to a single OCL or present in less than a quarter of all OCL types. Re-assessment of the annotations previously assigned to all genes based on the detection of protein domains/motifs and predicted functional roles [10, 13], revealed that some GtrOC proteins may either belong to new families or have been incorrectly assigned as glycosyltransferases. However, to date, there have been no studies conducted to confirm the specific roles of OCL genes in the construction of the OC. Therefore, annotations of genes at the OC locus may change in the future once experimental data on the function of GtrOC named proteins becomes available.

Across *Acinetobacter* spp., variation in gene content can also be observed just outside the OC locus, where a putative *waaL* O-antigen ligase gene can be found in some genomes, located between *aspS* and an unknown open reading frame that is immediately upstream of a gene coding for a TonB-dependent receptor [13]. However, in this study we did not detect any genes predicted to encode a WaaL ligase inserted at this location in the studied *A. baumannii* genomes. The absence of a *waaL* gene in the *A. baumannii* genome was first demonstrated in 2013 [10], and later confirmed using a wider variety of *A. baumannii* genome sequences in the following year [13]. While two *waaL*-like genes were later detected in some *A. baumannii* genomes [5, 29], these were later shown to encode ligases that link CPS oligosaccharide units to either the Type IV pilus or proteins through O-linked glycosylation [29, 30]. Absence of an O-antigen was also experimentally confirmed for isolate ATCC 17978 by the lack of an O-antigen banding pattern on Western blots using an anti-Lipid A antibody [14]. Hence, all available evidence to date provides strong support that *A. baumannii* produces LOS, with structural variation predominately seen in the OC.

Variation at the OC locus in clonal lineages has been reported extensively in previous studies, e.g. [7, 8, 10, 11, 13, 27, 28, 31–33]. In our previous large-scale analysis of >3600 genome sequences available at that time, we detected six OCL types in ST1 and four OCL types in both ST2 and ST25 [11]. Here, we report the finding of an additional four OCL in ST2, bringing the total to eight, with OCL1 being the most common. No additional OCL types in the ST1 lineage were found, though OCL1 also remains the most predominant type in this clone. ST25 was found to include 3 additional OCL but surprisingly does not carry OCL1. For the other over-represented STs, only one OCL type was detected in ST3, ST10, ST32 and ST78, two OCL in ST499 and four OCL in ST79. While we have detected additional variation at the OC locus in clonal isolates, as more genomes are sequenced and released, it is likely that further OCL forms will be observed amongst isolates belonging to the same ST. This highlights the importance of including OCL typing in epidemiological studies on *A. baumannii* clonal lineages.

## Acknowledgements

We thank Kelly Wyres from Monash University, Australia, and Kathryn Holt and Thomas Stanton from the London School of Hygiene & Tropical Medicine, UK, for their assistance with releasing the database and updated code on the *Kaptive* platforms.

