## Supplementary Figure S1 for "Identification of further variation at the lipooligosaccharide outer core locus in *Acinetobacter baumannii* genomes and extension of the OCL reference sequence database for *Kaptive*"

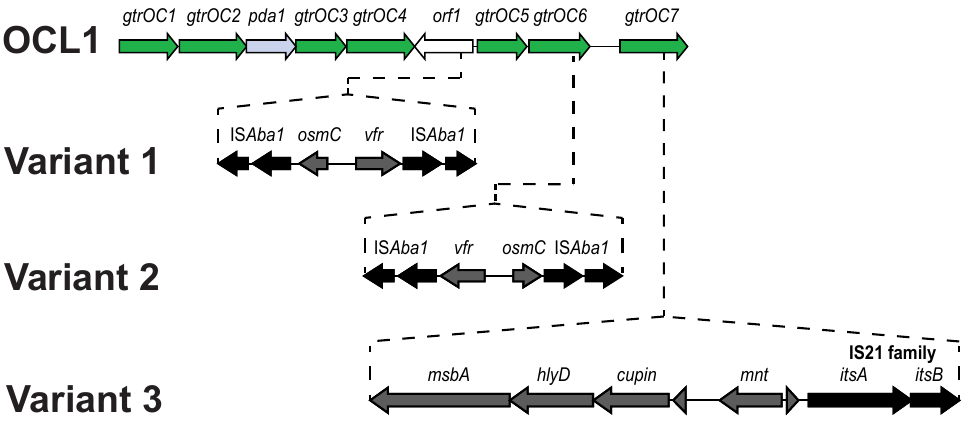

**Figure S1.** Transposon variants identified interrupting OCL1 indicating their specific insertion sites. Drawn to scale from NCBI accession numbers GCA_000516095.2 (variant 1), GCA_004101685.1 (variant 2), and GCA_021569095.1 (variant 3). Insertion sequences are black and other genes are grey. OCL1 colour scheme is shown in the legend of Figure 2.
