## Supplementary Figure S2 for "Identification of further variation at the lipooligosaccharide outer core locus in *Acinetobacter baumannii* genomes and extension of the OCL reference sequence database for *Kaptive*"

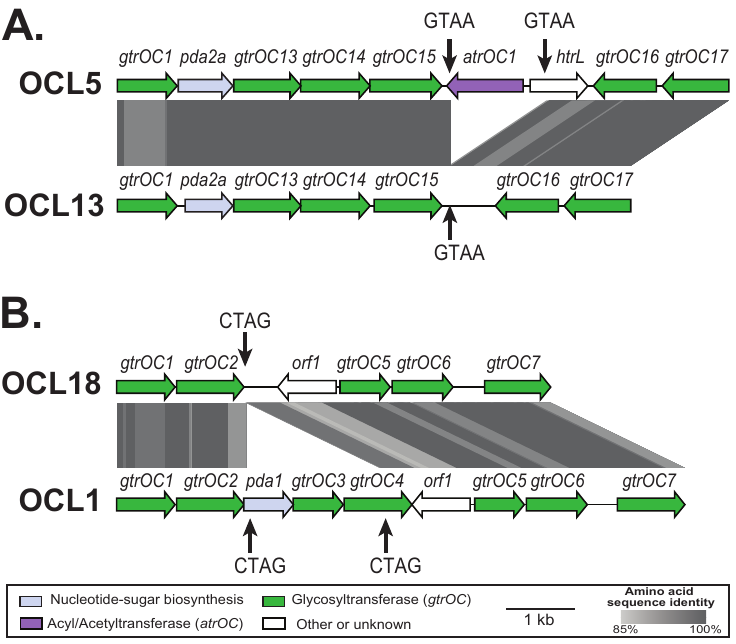

**Figure S2**. Comparison of *A. baumannii* OCL that include a 4 bp repeat influencing *Kaptive* assignments. **A**. Alignment of OCL5 and OCL13. **B.** Alignment of OCL18 and OCL1. Vertical black arrows indicate positions of the 4 bp repeats. Horizontal arrows are genes showing the direction of transcription, colour coded by the predicted function of their gene products. Grey shading between gene clusters shows amino acid sequence identity determined by tblastx. Colour scheme and grey scale are shown below. Figures drawn to scale using Easyfig [20] and annotated/coloured in Adobe Illustrator.
